# Oxidative stress response mediated by the yeast Rho5 GTPase depends on the proper spatiotemporal distribution of its dimeric GEF

**DOI:** 10.1101/2024.08.09.607359

**Authors:** Linnet Bischof, Jürgen J. Heinisch

**Affiliations:** Department of Biology/Chemistry, Division of Genetics, University of Osnabrück, Barbarastrasse 11, D-49076 Osnabrück, Germany

## Abstract

The small GTPase Rho5 acts as a central hub to mediate the yeast’s response to adverse environmental conditions, including oxidative stress, with the concomitant induction of mitophagy and apoptosis. A proper cellular stress response has been correlated with the rapid translocation of the GTPase to the mitochondria, which depends on its activating dimeric GDP/GTP exchange factor (GEF). Here, the small ALFA tag was attached to Rho5 or the GEF subunits Dck1 and Lmo1 to efficiently trap the functional fusion proteins to specific cellular membranes, i.e. the plasma membrane, the mitochondrial outer membrane, or the nuclear membrane, *via* fusions of integral membrane proteins residing in these compartments with an ALFA nanobody. The trapped components were subjected to life-cell fluorescence microscopy in combination with GFP fusions of the GTPase or its GEF subunits to investigate their interaction *in vivo*. We found that the dimeric GEF tends to auto-assemble and form stable dimers independent of its intracellular localization. On the other hand, GFP-Rho5 does not stably colocalize with the trapped GEF, attributed to its transient interaction. Phenotypic analyses of strains with the misslocalized proteins indicate that for a proper oxidative stress response Lmo1 needs to associate with the plasma membrane. In contrast, Rho5 only exerts its role at the mitochondrial surface when it is there in its active conformation. These data underline the importance of the proper spatio-temporal distribution of Rho5-GTP during oxidative stress response.

## 1. Introduction

GTPases comprise a huge family of molecular switches which mediate many cellular processes in eukaryotes, such as protein trafficking, cell motility, and signal transduction. Accordingly, their malfunctions have been associated with severe human conditions, such as diabetes, neurodegenerative diseases, and several forms of cancer (Crosas-Molist et al., 2022; Soriano et al., 2021; Stankiewicz and Linseman, 2014). For the subfamily of Ras-homologous GTPases (Rho-GTPases) six members have been identified in the yeast *Sacharomyces cerevisiae* (Rho1 to Rho5 and Cdc42), being involved in the regulation of cell wall synthesis (Rho1 and Rho5), and the establishment of cell polarity and bud site selection (Rho2 to Rho4 and Cdc42; reviewed in (Perez and Rincon, 2010)).

Rho5 was the last member of the subfamily to be identified in yeast, then as a negative regulator of the cell wall integrity (CWI) signaling pathway (Schmitz et al., 2002). As other GTPases, it is presumed to be active in its GTP-bound and inactive in its GDP-bound state. These states are interconverted by the help of a GDP/GTP exchange factor (GEF, suggested to be a dimer formed between Dck1 and Lmo1; (Schmitz et al., 2015)), and a GTPase activating protein (GAP), which promotes the intrinsic GTP hydrolyzing activity (Rgd2; (Annan et al., 2008)). In addition, GDP dissociation inhibitors (GDIs) enable the trafficking of Rho-GTPases through the cytosol in their inactive conformation by shielding the hydrophobic lipid anchor posttranslationaly attached to the cysteine residue of their CAAX box. However, Rdi1, the sole GDI found in *S. cerevisiae*, was shown to interact with Rho1, Rho4 and Cdc42, but not with Rho5 (Tiedje et al., 2008). Of note, Rho5 is the yeast homologue of human Rac1, with some interesting similarities in structure, subcellular distribution and physiological functions (reviewed in (Bischof et al., 2024a)). This includes the activation of Rac1 by dimeric GEFs of the DOCK/ELMO family, after which the yeast GEF subunits have been named. Homologues of Rho5/Rac1 are also found in other yeast and filamentous fungi, with functions in the establishment of cell polarity, hyphal growth, pathogenicity, or dealing with reactive oxygen species (reviewed in (Hühn et al., 2020)).

In *S. cerevisiae*, besides the role in CWI signaling mentioned above, Rho5 was shown to also affect the high osmolarity glycerol (HOG) signal transduction pathway (Annan et al., 2008), the response to oxidative stress (Singh et al., 2008), and the reaction to glucose starvation (Schmitz et al., 2018; Schweitzer et al., 2024). Both the exposure to hydrogen peroxide and the depletion of glucose have been shown to trigger the translocation of Rho5 from the yeast plasma membrane to the mitochondrial surface (Schmitz et al., 2018; Schmitz et al., 2015). This has been proposed to activate mitophagy and apoptosis, at least in response to oxidative stress (Schmitz et al., 2015). In the latter study we also showed that the GTPase only translocates in cells with an active GEF, being drastically reduced in *dck1* or *lmo1* deletions. The GEF subunits also translocate to mitochondria under oxidative stress, but independent from each other and from the presence of Rho5 (Bischof et al., 2022). Several proteins of the mitochondrial outer membrane, such as Por1 and Msp1, have been proposed to participate in the recruitment of Rho5 to mitochondria under stress, but not of Dck1 and Lmo1, further substantiating that the GTPase and its GEF translocate independent from each other and possibly by using different mechanisms (Bischof et al., 2024b).

Despite these investigations, the question of how and in which order the three proteins associate *in vivo* and where in the cell the activated Rho5 fulfils its physiological functions has not yet been sufficiently addressed. In this work, we used fusion constructs with different transmembrane proteins to direct the highly specific ALFA nanobody to specific cellular membranes. The ALFA-tagged GTPase or its GEF subunits could thus be efficiently recruited to these sites to study the physiological consequences of their misslocalization.

## 2. Results

### 2.1. The GEF subunits Lmo1 and Dck1 auto-assemble in vivo independent of their association with different endomembranes

To address the question in which order and under which conditions Rho5 and its GEF associate within the yeast cell, we assumed that the dimeric GEF may be formed first in a fairly stable, closed conformation, as observed for the human DOCK/ELMO complex, whose structure has been elucidated (Chang et al., 2020; Kukimoto-Niino et al., 2024). An ALFA tag was therefore attached to the N-terminal end of Lmo1 (ALFA-Lmo1) and the C-terminal end of Dck1 (Dck1-ALFA), using a PAPAP linker previously proven to retain maximal functionality of the respective proteins with GFP fusions (Bischof et al., 2022). To avoid variations in copy numbers and gene expression, all fusion proteins employed in this work were either encoded at the native genetic locus or integrated at the *ura3-52* locus and expressed from their native promoters in otherwise isogenic strains (listed in Table 1). They were then combined by conventional crossing and tetrad analysis for the following investigations.

**Table 1.** Yeast strains employed in this work.

| Designation | Genotype <sup>a,b</sup> | Reference |
| --- | --- | --- |
| FSO23-3B | *MATalpha dck1::KILEU2* | this work |
| FSO23-6B | *MATa dck1::KILEU2* | this work |
| HD56/AnbH11 | *MATalpha NUP159-ALFAnb-SpHIS5* | this work |
| HD56-5A | *MATalpha* | (Arvanitidis and Heinisch, 1994) |
| HLBO105-1B | *MATa NUP159-ALFAnb-SpHIS5* | this work |
| HLBO106n-14C | *MATalpha ALFA-LMO1-KILEU2 NUP159-ALFAnb-SpHIS5 DCK1-3myEGFP-SpHIS5* | this work |
| HLBO106n-1C | *MATalpha ALFA-LMO1-KILEU2 NUP159-ALFAnb-SpHIS5* | this work |
| HLBO106n-2A | *MATalpha DCK1-3myEGFP-SpHIS5 ALFA-LMO1-KILEU2 IDP1-mCherry-kanMX NUP159-ALFAnb-SpHIS5* | this work |
| HLBO106n-7A | *MATalpha DCK1-3myEGFP-SpHIS5 ALFA-LMO1-KILEU2 IDP1-mCherry-kanMX NUP159-ALFAnb-SpHIS5* | this work |
| HLBO111-1C | *MATalpha DCK1-ALFATag-SpHIS5 GFP-LMO1-KILEU2* | this work |
| HLBO111-6D | *MATa IDP1-mCherry-kanMX DCK1-ALFATag-SpHIS5 GFP-LMO1-KILEU2 NUP159-ALFAnb-SpHIS5* | this work |
| HLBO111-8C | *MATalpha DCK1-ALFATag-SpHIS5 NUP159-ALFAnb-SpHIS5* | this work |
| HLBO112-4C | *MATa ALFA-LMO1-KILEU2 IDP1-mCherry-kanMX GFP-pl-RHO5-SkHIS3 NUP159-ALFAnb-SpHIS5* | this work |
| HLBO112-6B | *MATa ALFA-LMO1-KILEU2 IDP1-mCherry-kanMX GFP-pl-RHO5-SkHIS3 NUP159-ALFAnb-SpHIS5* | this work |
| HLBO113.2-2A | *MATalpha ALFA-LMO1-KILEU2 IDP1-mCherry-kanMX GFP-pl-RHO5-SkHIS3 MID2-ALFAnb-kanMX* | this work |
| HLBO114-2C | *MATa ura3-52::URA3-ALFAnb-FIS1<sup>TMD</sup> ALFA-LMO1-KILEU2 IDP1-mCherry-kanMX GFP-pl-RHO5-SkHIS3* | this work |
| HLBO114-9C | *MATalpha ura3-52::URA3-ALFAnb-FIS1<sup>TMD</sup> ALFA-LMO1-KILEU2 IDP1-mCherry-kanMX GFP-RHO5::SkHIS3 linker3* | this work |
| HLBO115-3A | *MATalpha IDP1-mCherry-kanMX ura3-52::URA3-ALFAnb-FIS1<sup>TMD</sup> GFP-LMO1-KILEU2 ALFA-RHO5::SkHIS3* | this work |
| HLBO115-8C | *MATa IDP1-mCherry-kanMX ura3-52::URA3-ALFAnb-FIS1<sup>TMD</sup> GFP-LMO1-KILEU2 ALFA-RHO5::SkHIS3* | this work |
| HLBO116-11A | *MATalpha ALFA-RHO5-SkHIS3 IDP1-mCherry-kanMX GFP-LMO1-KILEU2 NUP159-ALFAnb-SpHIS5* | this work |
| HLBO116-2B | *MATalpha ALFA-RHO5-SkHIS3 IDP1-mCherry-kanMX GFP-LMO1-KILEU2 NUP159-ALFAnb-SpHIS5* | this work |
| HLBO117-1A | *MATalpha ALFA-RHO5-SkHIS3 NUP159-ALFAnb-SpHIS5* | this work |
| HLBO117-1C | *MATa ALFA-RHO5-SkHIS3 NUP159-ALFAnb-SpHIS5* | this work |
| HLBO117-6B | *MATalpha ALFA-RHO5-SkHIS3 NUP159-ALFAnb-SpHIS5* | this work |
| HLBO117-6D | *MATa ALFA-RHO5-SkHIS3 NUP159-ALFAnb-SpHIS5* | this work |
| HLBO20-4B | *MATa* | this work |
| HLBO21-10A | *MATalpha DCK1-3GFP-SpHIS5 IDP1-mCherry-kanMX* | this work |
| HLBO21-2A | *MATa DCK1-3GFP-SpHIS5 IDP1-mCherry-kanMX* | this work |
| HLBO27-5D | *MATa IDP1-mCherry-SkHIS3 GFP-LMO1-KILEU2* | this work |
| HLBO31-3B | *MATa IDP1-mCherry-SkHIS3 GFP-LMO1-KILEU2* | this work |
| HLBO38-1A | *MATalpha ura3-52::URA3-ALFAnb-FIS1<sup>TMD</sup>* | this work |
| HLBO38-2B | *MATa ura3-52::URA3-ALFAnb-FIS1<sup>TMD</sup> GFP-LMO1-KILEU2* | this work |
| HLBO38-8A | *MATalpha* | this work |
| HLBO38-8D | *MATa DCK1-ALFATag-SpHIS5* | this work |
| HLBO38-9A | *MATa ura3-52::URA3-ALFAnb-FIS1<sup>TMD</sup> DCK1-ALFATag-SpHIS5* | this work |
| HLBO39-10B | *MATalpha ura3-52::URA3-ALFAnb-FIS1<sup>TMD</sup> DCK1-ALFATag-SpHIS5* | this work |
| HLBO39-1B | *MATalpha DCK1-ALFATag-SpHIS5* | this work |
| HLBO39-1B- | *MATalpha DCK1-ALFATag-SpHIS5* | this work |
| HLBO39-3B | *MATalpha ura3-52::URA3-ALFAnb-FIS1<sup>TMD</sup> GFP-LMO1-KILEU2* | this work |
| HLBO43-13C | *MATa IDP1-mCherry::kanMX ura3-52::URA3-ALFAnb-FIS1<sup>TMD</sup> DCK1-ALFATag-SpHIS5 GFP-LMO1-KILEU2* | this work |
| HLBO44-10A | *MATalpha IDP1-mCherry-kanMX ura3-52::URA3-ALFAnb-FIS1<sup>TMD</sup> DCK1-ALFATag-SpHIS5 GFP-LMO1-KILEU2* | this work |
| HLBO56-10A | *MATalpha DCK1-3myEGFP-SpHIS5 ALFA-LMO1-KILEU2 IDP1-mCherry-kanMX ura3-52::URA3-ALFAnb-FIS1<sup>TMD</sup>* | this work |
| HLBO56-11D | *MATa ALFA-LMO1-KILEU2 ura3-52::URA3-ALFAnb-FIS1<sup>TMD</sup>* | this work |
| HLBO56-2A | *MATa DCK1-3myEGFP-SpHIS5 ALFA-LMO1-KILEU2 IDP1-mCherry-kanMX ura3-52::URA3-ALFAnb-FIS1<sup>TMD</sup>* | this work |
| HLBO56-2D | *MATa ALFA-LMO1-KILEU2 ura3-52::URA3-ALFAnb-FIS1<sup>TMD</sup>* | this work |
| HLBO56-4B | *MATa DCK1-3myEGFP-SpHIS5 ALFA-LMO1-KILEU2 IDP1-mCherry-kanMX ura3-52::URA3-ALFAnb-FIS1<sup>TMD</sup>* | this work |
| HLBO56-5A | *MATalpha ALFA-LMO1-KILEU2 ura3-52::URA3-ALFAnb-FIS1<sup>TMD</sup>* | this work |
| HLBO57-5A | *MATalpha ALFA-LMO1-KILEU2 ura3-52::URA3-ALFAnb-FIS1<sup>TMD</sup>* | this work |
| HLBO60-11C | *MATalpha ALFA-LMO1-KILEU2 IDP1-mCherry-kanMX GFP-pl-RHO5-SkHIS3* | this work |
| HLBO60-7D | *MATa IDP1-mCherry-kanMX GFP-pl-RHO5-SkHIS3* | this work |
| HLBO61-1A | *MATa IDP1-mCherry-kanMX GFP-pl-RHO5-SkHIS3* | this work |
| HLBO83-1A | *MATa ura3-52::URA3-ALFAnb-FIS1<sup>TMD</sup>* | this work |
| HLBO83-1B | *MATalpha ura3-52::URA3-ALFAnb-FIS1<sup>TMD</sup> ALFA-RHO5-SkHIS3* | this work |
| HLBO83-2A | *MATa ura3-52::URA3-ALFAnb-FIS1<sup>TMD</sup> ALFA-RHO5-SkHIS3* | this work |
| HLBO83-4A | *MATa ura3-52::URA3-ALFAnb-FIS1<sup>TMD</sup> ALFA-RHO5-SkHIS3* | this work |
| HLBO83-5D | *MATalpha ura3-52::URA3-ALFAnb-FIS1<sup>TMD</sup>* | this work |
| HLBO83-7D | *MATa ALFA-RHO5-SkHIS3* | this work |
| HLBO83-7D | *MATa ALFA-RHO5-SkHIS3* | this work |
| HLBO84-10D | *MATalpha ALFA-LMO1-KILEU2 MID2-ALFAnb-kanMX* | this work |
| HLBO84-1D | *MATalpha DCK1-3myEGFP-SpHIS5 ALFA-LMO1-KILEU2 IDP1-mCherry-kanMX MID2-ALFAnb-kanMX* | this work |
| HLBO84-2B | *MATalpha ALFA-LMO1-KILEU2 MID2-ALFAnb-kanMX* | this work |
| HLBO84-7D | *MATa DCK1-3myEGFP-SpHIS5 ALFA-LMO1-KILEU2 IDP1-mCherry-kanMX MID2-ALFAnb-kanMX* | this work |
| HLBO85-1A | *MATa ALFA-RHO5-SkHIS3 MID2-ALFAnb-kanMX* | this work |
| HLBO85-1D | *MATalpha MID2-ALFAnb-kanMX* | this work |
| HLBO85-1D | *MATalpha MID2-ALFAnb-kanMX* | this work |
| HLBO85-2C | *MATalpha MID2-ALFAnb-kanMX* | this work |
| HLBO85-2C | *MATalpha MID2-ALFAnb-kanMX* | this work |
| HLBO85-3B | *MATalpha ALFA-RHO5-SkHIS3 MID2-ALFAnb-kanMX* | this work |
| HLBO85-4C | *MATalpha MID2-ALFAnb-kanMX* | this work |
| HLBO85-6C | *MATa ALFA-RHO5-SkHIS3 MID2-ALFAnb-kanMX* | this work |
| HLBO85-7C | *MATalpha ALFA-RHO5-SkHIS3* | this work |
| HLBO85-7C | *MATalpha ALFA-RHO5-SkHIS3* | this work |
| HLBO86-4B | *MATalpha ALFA-RHO5-SkHIS3* | this work |
| HLBO88-10D | *MATalpha MID2-ALFAnb-kanMX DCK1-ALFATag-SpHIS5* | this work |
| HLBO88-12A | *MATalpha MID2-ALFAnb-kanMX DCK1-ALFATag-SpHIS5* | this work |
| HLBO88-4C | *MATalpha MID2-ALFAnb-kanMX IDP1-mCherry-kanMX DCK1-ALFATag-SpHIS5 GFP-LMO1-KILEU2* | this work |
| HLBO88-8A | *MATa MID2-ALFAnb-kanMX IDP1-mCherry-kanMX DCK1-ALFATag-SpHIS5 GFP-LMO1-KILEU2* | this work |
| HLBO89-1D | *MATalpha DCK1-ALFATag-SpHIS5 GFP-LMO1-KILEU2 NUP159-ALFAnb-SpHIS5* | this work |
| HLBO89-2C | *MATa DCK1-ALFATag-SpHIS5* | this work |
| HLBO91n-12D | *MATalpha ALFA-RHO5-SkHIS3 MID2-ALFAnb-kanMX IDP1-mCherry-kanMX GFP-LMO1-KILEU2* | this work |
| HLBO91n-13B | *MATalpha ALFA-RHO5-SkHIS3 MID2-ALFAnb-kanMX IDP1-mCherry-kanMX GFP-LMO1-KILEU2* | this work |
| HOD464-7B | *MATa lmo1::kanMX* | this work |
| HOD505-1A | *MATalpha dck1::KILEU2* | this work |
| HOD505-1C | *MATa dck1::KILEU2* | this work |
| HOD555-1A | *MATa* | (Bischof et al., 2024b) |
| HOD555-1D | *MATa rho5::kanMX* | (Bischof et al., 2024b) |
| HOD555-3D | *MATa rho5::kanMX* | (Bischof et al., 2024b) |
| HOD556-2B | *MATalpha* | (Bischof et al., 2024b) |
| LBO309 | *MATalpha ALFA-LMO1-KILEU2* | this work |
| LBO310 | *MATa ALFA-LMO1-KILEU2* | this work |
| LBO81 | *MATa lmo1::KIURA3* | this work |

<sup>a</sup> All strains used in this work were derived from the isogenic diploid strain DHD5 (Kirchrath et al., 2000) and shared the genotype *his3-11,15 leu2-3112 ura3-52 MAL3 SUC2 GAL*. Therefore, only the mating type and the loci manipulated for this study are listed.
<sup>b</sup> The constructs with *ura3-52::URA3-ALFAnb-FIS1<sup>TMD</sup>* were obtained by linearizing the yeast integrative plasmid carrying the fusion construct with EcoRV prior to transformation of the recipient strain and selection for uracil prototrophy.

First, strains producing ALFA-tagged Lmo1 were individually combined with those containing ALFA nanobody fusions at either the mitochondrial surface (ALFAnb-Fis1^TMD^), the plasma membrane (Mid2-ALFAnb), or the nuclear surface (Nup159-ALFAnb). Furthermore, segregants also carrying a *DCK1-GFP* allele were obtained from different crosses, to follow the intracellular distribution of the encoded GEF subunit fusion by life-cell fluorescence microscopy. As evident from Fig. 1A, Dck1-GFP displayed a diffused punctate pattern in non-stressed cells devoid of an ALFA nanobody construct to trap Lmo1 to any specific membrane. Addition of hydrogen peroxide then led to the rapid translocation of Dck1-GFP to mitochondria, as would be expected from previous results (Bischof et al., 2022; Schmitz et al., 2015). In contrast, strains in which ALFA-Lmo1 was confined to the mitochondrial surface also showed the Dck1-GFP signal at these organelles, no matter if the cells were grown under standard conditions or stressed with hydrogen peroxide. Likewise, ALFA-Lmo1 trapped at the plasma membrane or at the nuclear membrane by the appropriate nanobody fusions recruited Dck1-GFP to the respective sites in the absence or presence of hydrogen peroxide, as judged from the processed microscopy images (Fig. 1A). Association of the GFP signal to mitochondria under stress was reduced by at least twofold when Lmo1 was trapped at the plasma membrane, and more than tenfold upon its association with the nuclear surface (Fig. 1B). We conclude that Dck1 efficiently interacts with Lmo1 independent of the intracellular compartment the latter resides at, and probably assembles into a GEF dimer.

**Figure 1:**
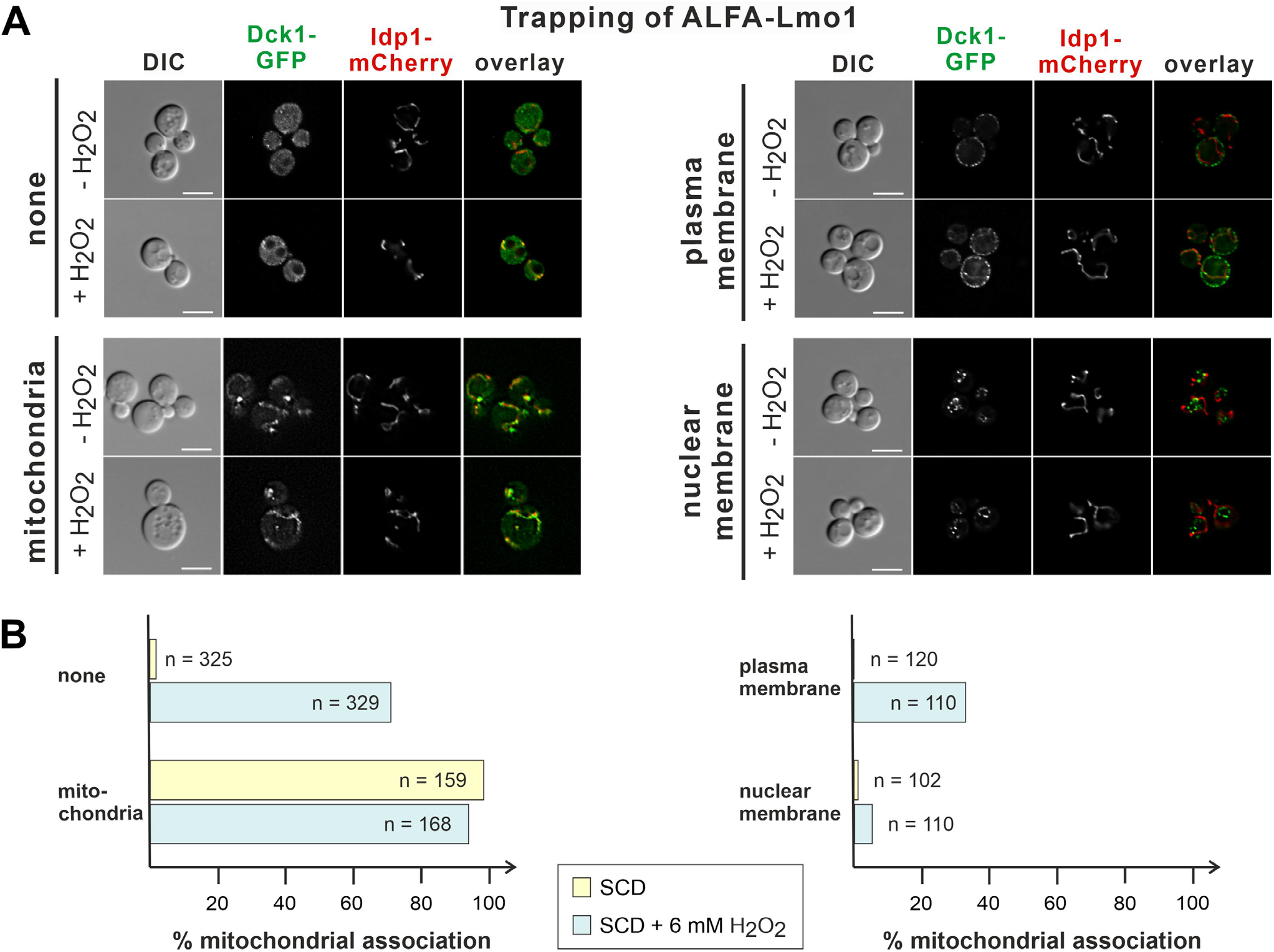
ALFA-Lmo1 trapped at different subcellular compartments recruits Dck1-GFP. The distribution of Dck1-GFP was followed in strains where ALFA-Lmo1 is trapped at either the mitochondrial outer membrane (ALFAnb-Fis1^TMD^), the plasma membrane (Mid2-ALFAnb) or the nucleus (Nup159-ALFAnb). A) A representative set of cells with the indicated combinations of traps and fluorescently labelled proteins is shown. To obtain the microscopic images, cells were grown on synthetic medium with 2% glucose. Where indicated, cells were exposed for 5 to 10 minutes to 6 mM hydrogen peroxide prior to image acquisition. For visualization of mitochondria, Idp1-mCherry was used, expressed from a gene fusion at the native locus. Size bars in the DIC (differential interference contrast) images correspond to 5 μm and are applicable to all images in the same row. B) Quantification of the results from life-cell fluorescence microscopy. Percentages of cells showing the Dck1-GFP signal associated with mitochondria were calculated from the total cell counts (n), with or without hydrogen peroxide as indicated. At least two independent isogenic segregants were examined for each data set. Strains employed (see Table 1 for genotypes): Dck1-GFP (HLBO21-2A and HLBO21-10A), Fis1-Lmo1 trap Dck1-GFP (HLBO56-2A, HLBO56-4B and HLBO56-10A), Mid2-Lmo1 trap Dck1-GFP (HLBO84-1D and HLBO84-7D), Nup159-Lmo1 trap Dck1-GFP (HLBO106n-2A and HLBO106n-7A).

To substantiate this notion, a reverse approach was followed. Thus, strains with the ALFA-tagged Dck1 subunit were combined with the ALFA nanobodies confined to the different membranes, following the distribution of GFP-Lmo1 (Fig. 2A). Again, under standard growth conditions Lmo1 was recruited to the sites where Dck1 was trapped, indicating the formation of the dimeric GEF complex. Under oxidative stress, the translocation of GFP-Lmo1 to mitochondria was reduced more than twofold in strains confining Dck1 to the plasma membrane or to the nuclear surface, as compared to the strain not carrying any membrane-trap (Fig. 2B). While these results suggest that the GEF subunits can auto-assemble *in vivo* independent of their subcellular localization, we noted that Lmo1 seems to have a higher capacity of retaining Dck1 at the nuclear surface under oxidative stress than *vice versa*. Therefore, trapping of ALFA-Lmo1 at a membrane was generally employed in the following experiments to direct the localization of the dimeric GEF.

**Figure 2:**
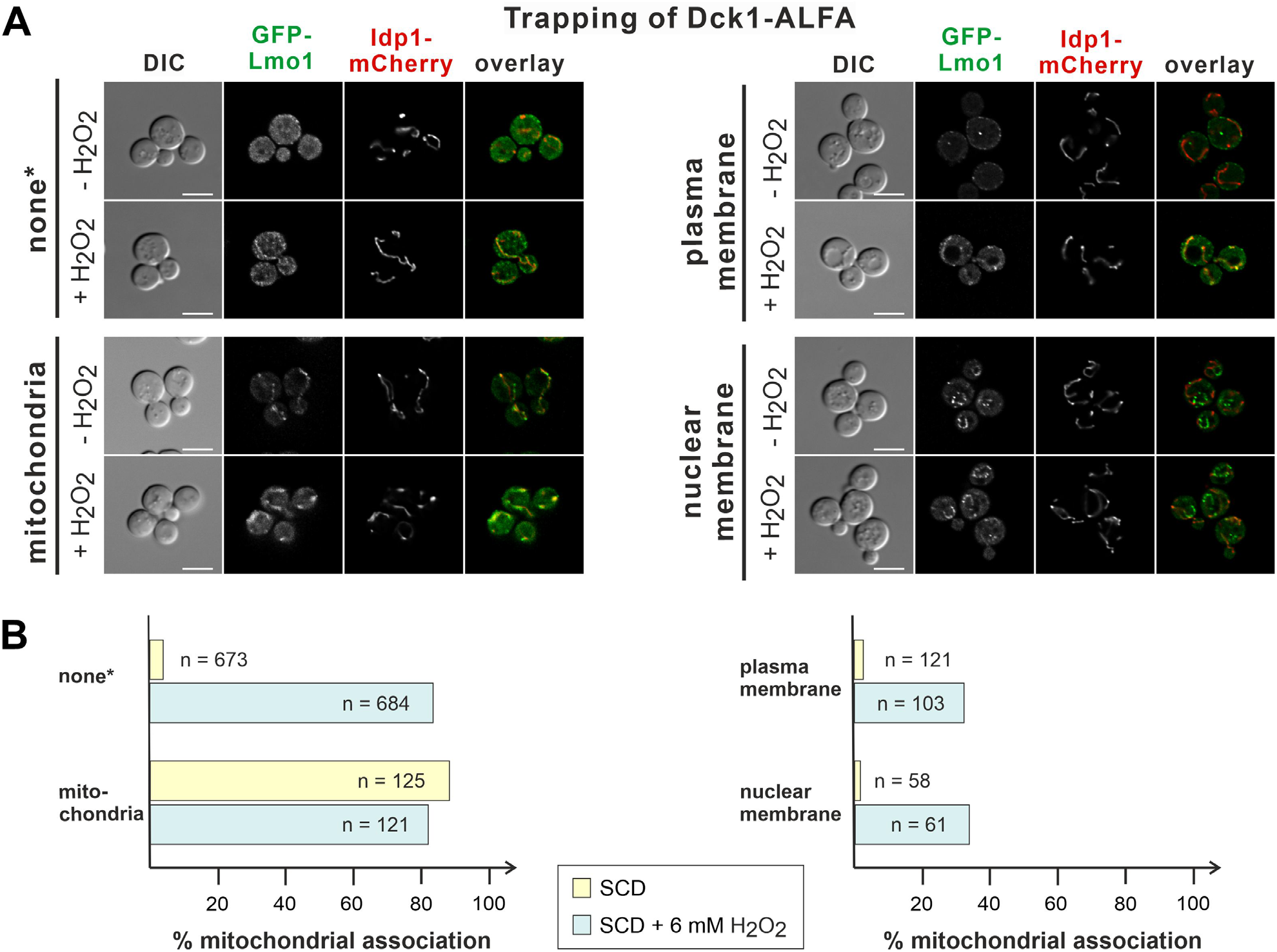
Dck1-ALFA trapped at different subcellular compartments recruits GFP-Lmo1. Dck1-ALFA was trapped by ALFA nanobody fusions at the compartments indicated as explained in the legend of Fig. 1. The strains also contained GFP-Lmo1 expressed from a gene fusion construct at the native *LMO1* locus. A) A representative set of cells with the indicated combinations of traps and fluorescently labelled proteins is shown. Growth conditions, exposure to hydrogen peroxide, and image acquisition were as outlined in the legend of Fig. 1. Size bars are 5 μm, applicable to all images in the same row. B) Quantification of the results from life-cell fluorescence microscopy. Percentages of cells showing the GFP-Lmo1 signal associated with mitochondria were calculated from the total cell counts (n), with or without hydrogen peroxide as indicated. At least two independent isogenic segregants were examined for each data set, with the exception of the nuclear trapping device, for which only one segregant was used for imaging. Strains employed (see Table 1 for genotypes): Fis1-Dck1 trap GFP-Lmo1 (HLBO43-13C and HLBO44-10A), Mid2-Dck1 trap GFP-Lmo1 (HLBO88-4C and HLBO88-8A), Nup159-Dck1 trap GFP-Lmo1 (HLBO111-6D). **\*** Note that for the control strains without a trap images in A) and cell counts in B) are the same here and in Fig. 4, where they have been inserted again for the sake of comparison.

### 2.2. Rho5 translocates to the mitochondrial surface independent from the localization of its GEF

From the data presented above it can be assumed that the trapped ALFA-Lmo1 forms a functional GEF with the untagged Dck1 at the respective membranes. It has been proposed that other GEFs associate with their target membranes and then recruit their GTPases for activation (reviewed in (Wright et al., 2014). Therefore, we next asked whether Rho5 can be recruited to the membranes that ALFA-Lmo1/Dck1 are associated with. For this purpose, the strains with the ALFA-Lmo1 traps were combined with strains producing a Rho5-GFP fusion protein (Fig. 3A).

**Figure 3:**
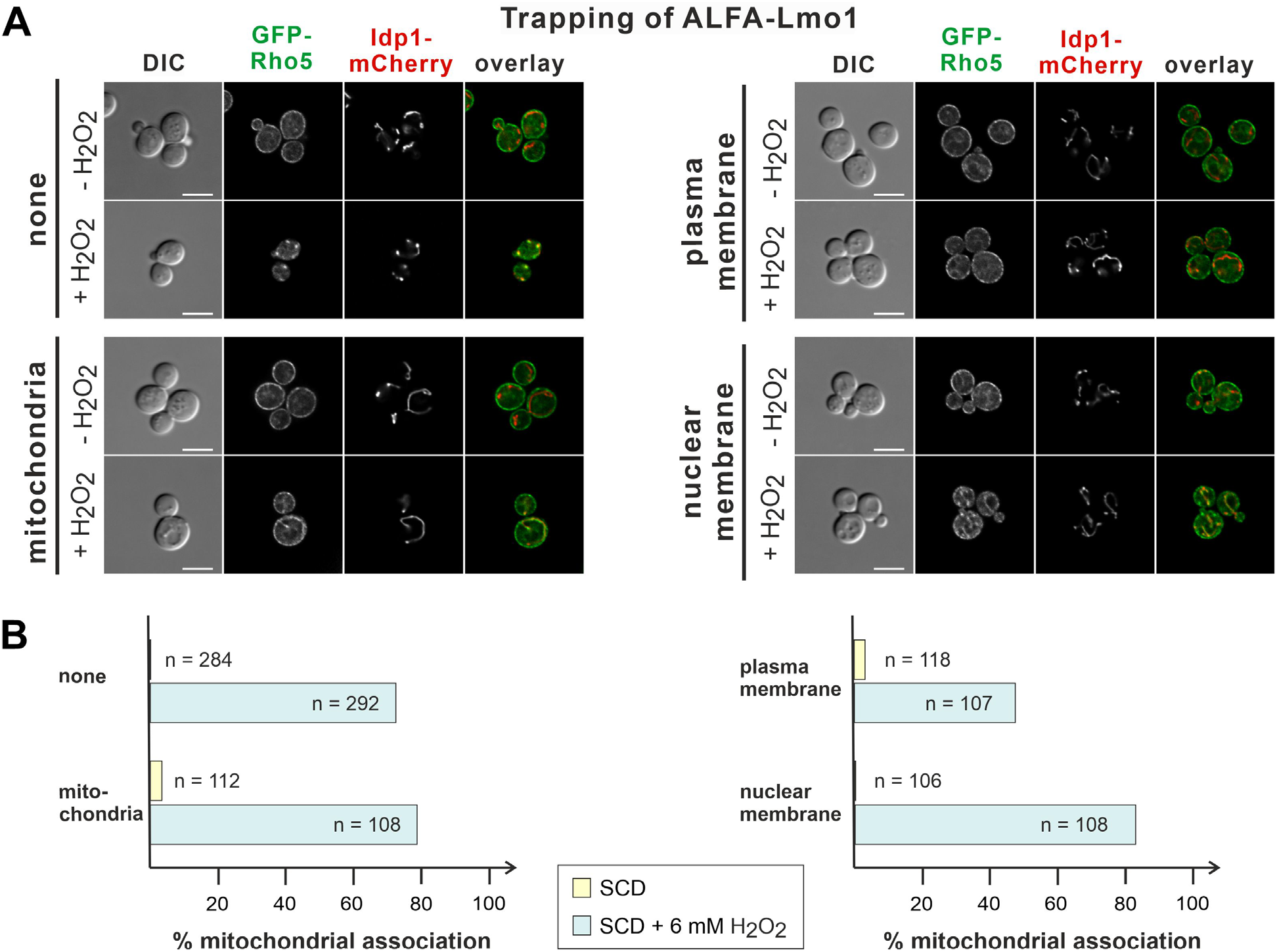
The intracellular distribution of GFP-Rho5 is barely affected by confining its GEF to different sub-cellular compartments. ALFA-Lmo1 was trapped by ALFA nanobody fusions at the compartments indicated as explained in the legend of Fig. 1. The strains also contained GFP-Rho5 expressed from a gene fusion construct at the native *RHO5* locus. A) A representative set of cells with the indicated combinations of traps and fluorescently labelled proteins is shown. Growth conditions, exposure to hydrogen peroxide, and image acquisition were as outlined in the legend of Fig. 1. Size bars are 5 μm, applicable to all images in the same row. B) Quantification of the results from life-cell fluorescence microscopy. Percentages of cells showing the GFP-Rho5 signal associated with mitochondria were calculated from the total cell counts (n), with or without hydrogen peroxide as indicated. At least two independent isogenic segregants were examined for each data set, with the exception of the plasma membrane trapping device, for which only one segregant was used for imaging. Strains employed (see Table 1 for genotypes): GFP-Rho5 (HLBO60-7D and HLBO61-1A), Fis1-Lmo1 trap GFP-Rho5 (HLBO114-2C and HLBO114-9C), Mid2-Lmo1 trap GFP-Rho5 (HLBO113.2-2A only one), Nup159-Lmo1 trap GFP-Rho5 (HLBO112-4C and HLBO112-6B).

In contrast to Dck1-GFP, the GTPase mostly appeared at the plasma membrane in non-stressed cells, independent of the localization of the tagged Lmo1. Addition of hydrogen peroxide invariably led to the translocation of GFP-Rho5 to mitochondria, which was only notably reduced by approximately 33% in strains that had Lmo1 trapped at the plasma membrane, as compared to the control without an ALFA nanobody fusion (Fig. 3B). This suggests that the interaction of the GEF with its GTPase is very transient and allows for free trafficking of the latter, as would be expected.

The latter view is supported by the reverse experiment, in which an ALFA-tagged Rho5 was trapped at the plasma membrane by the Mid2-ALFAnb fusion and monitoring the distribution of GFP-Lmo1 (Fig. 4A). Under oxidative stress, GFP-Lmo1 translocated in less than 10% of the cells examined to mitochondria as compared to the control strain without an ALFA nanobody fusion (Fig. 4B).

**Figure 4:**
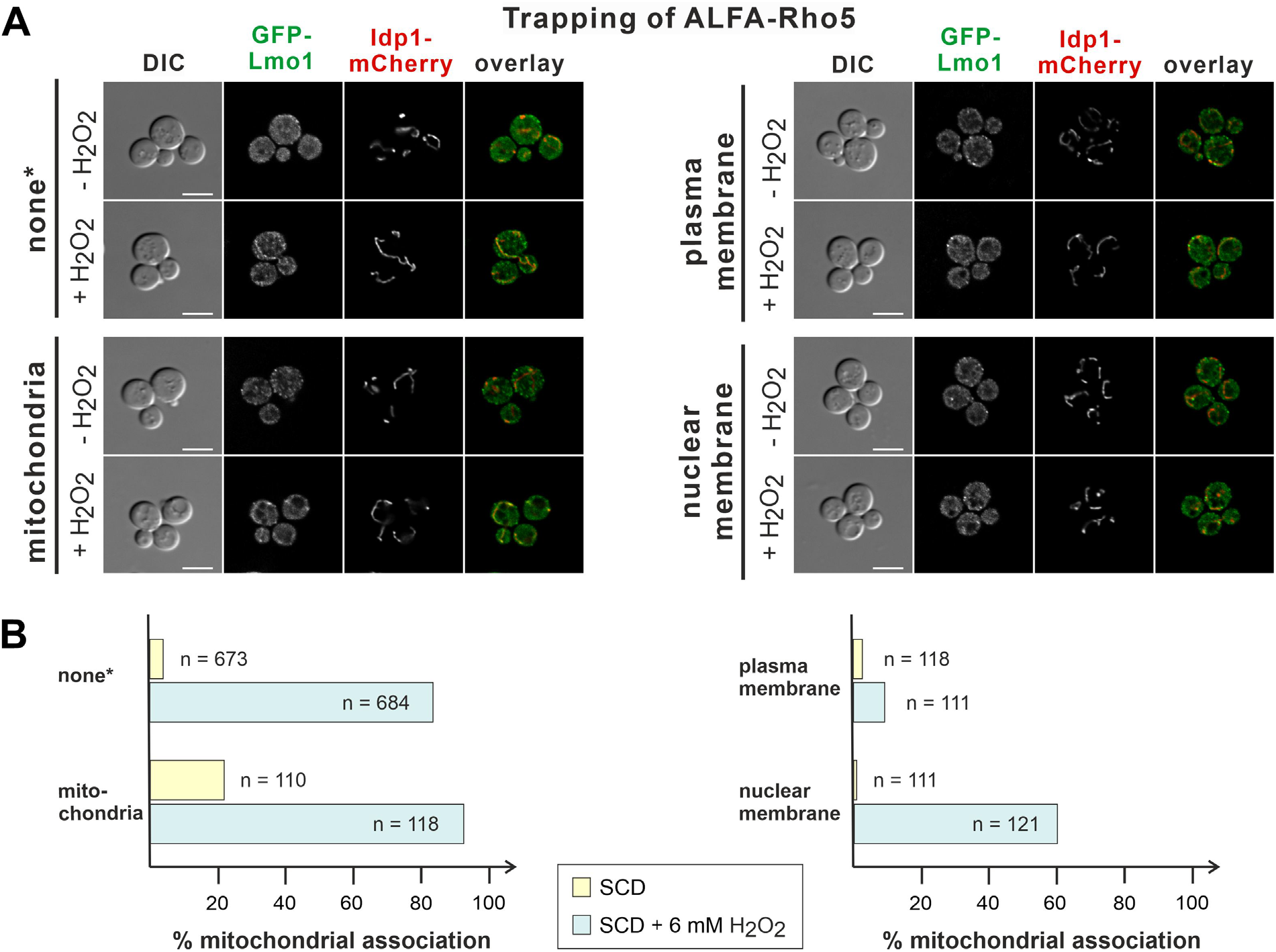
ALFA-Rho5 trapped at different subcellular compartments does not recruit GFP-Lmo1. ALFA-Rho5 was trapped by ALFA nanobody fusions at the compartments indicated as explained in the legend of Fig. 1. The strains also contained GFP-Lmo1 expressed from a gene fusion construct at the native *LMO1* locus. A) A representative set of cells with the indicated combinations of traps and fluorescently labelled proteins is shown. Growth conditions, exposure to hydrogen peroxide, and image acquisition were as outlined in the legend of Fig. 1. Size bars are 5 μm, applicable to all images in the same row. B) Quantification of the results from life-cell fluorescence microscopy. Percentages of cells showing the GFP-Lmo1 signal associated with mitochondria were calculated from the total cell counts (n), with or without hydrogen peroxide as indicated. At least two independent isogenic segregants were examined for each data set. Strains employed (see Table 1 for genotypes): Fis1-Rho5 trap GFP-Lmo1 (HLBO115-3A and HLBO115-8C), Mid2-Rho5 trap GFP-Lmo1 (HLBO91n-12D and HLBO91n-13B), Nup159-Rho5 trap GFP-Lmo1 (HLBO116-2B and HLBO116-11A). **\*** Note that for the control strains without a trap images in A) and cell counts in B) are the same as in Fig. 2. They have been inserted here again for the sake of comparison.

Tethering of Rho5 to the mitochondrial outer membrane resulted in a more than fivefold increase in the number of cells showing the GFP-Lmo1 signal at these organelles as compared to the control under standard growth conditions, whereas confining the GTPase to the nuclear envelope reduced the translocation of the GEF subunit to mitochondria under oxidative stress only by approximately 27% (Fig. 4B). A clear nuclear localization of GFP-Lmo1 could not be observed in this strain, neither in stressed, nor in non-stressed cells (Fig. 4A).

### 2.3. Confining Rho5 or its GEF subunits to specific endomembranes affects the yeast’s capacity to cope with oxidative stress

In order to determine the importance of intracellular distribution of Rho5 and its GEF subunits for the oxidative stress response, the strains examined in life-cell fluorescence microscopy were then employed to determine the phenotypes associated with confining each of the three proteins to a specific membrane. For this purpose, growth was followed in synthetic complete medium with 2% glucose in the presence and abscence of hydrogen peroxide (Fig. 5 and supplementary Fig. S1). A *rho5* deletion is clearly hyper-resistant towards oxidative stress, confirming previous observations (Schmitz et al., 2015; Sterk et al., 2019), whereas segregants with ALFA-Rho5 directed to the mitochondria by the Fis1-nanobody fusion grew similar to the wild-type strains (Fig. 5A). This suggests that Rho5 can fullfil its function in oxidative stress response when it resides at the mitochondrial surface. In contrast, directing the GTPase either to the plasma membrane or to the nuclear surface rendered the cells clearly hyper-resistant, even exceeding that of the deletion mutant (Figs. 5B and C; note that the increased resistance of the strains with only the ALFA-tagged Rho5, but not the nanobody construct, was not observed in the growth curves recorded for Fig. 5A and in many more growth assays not included in this work).

**Figure 5:**
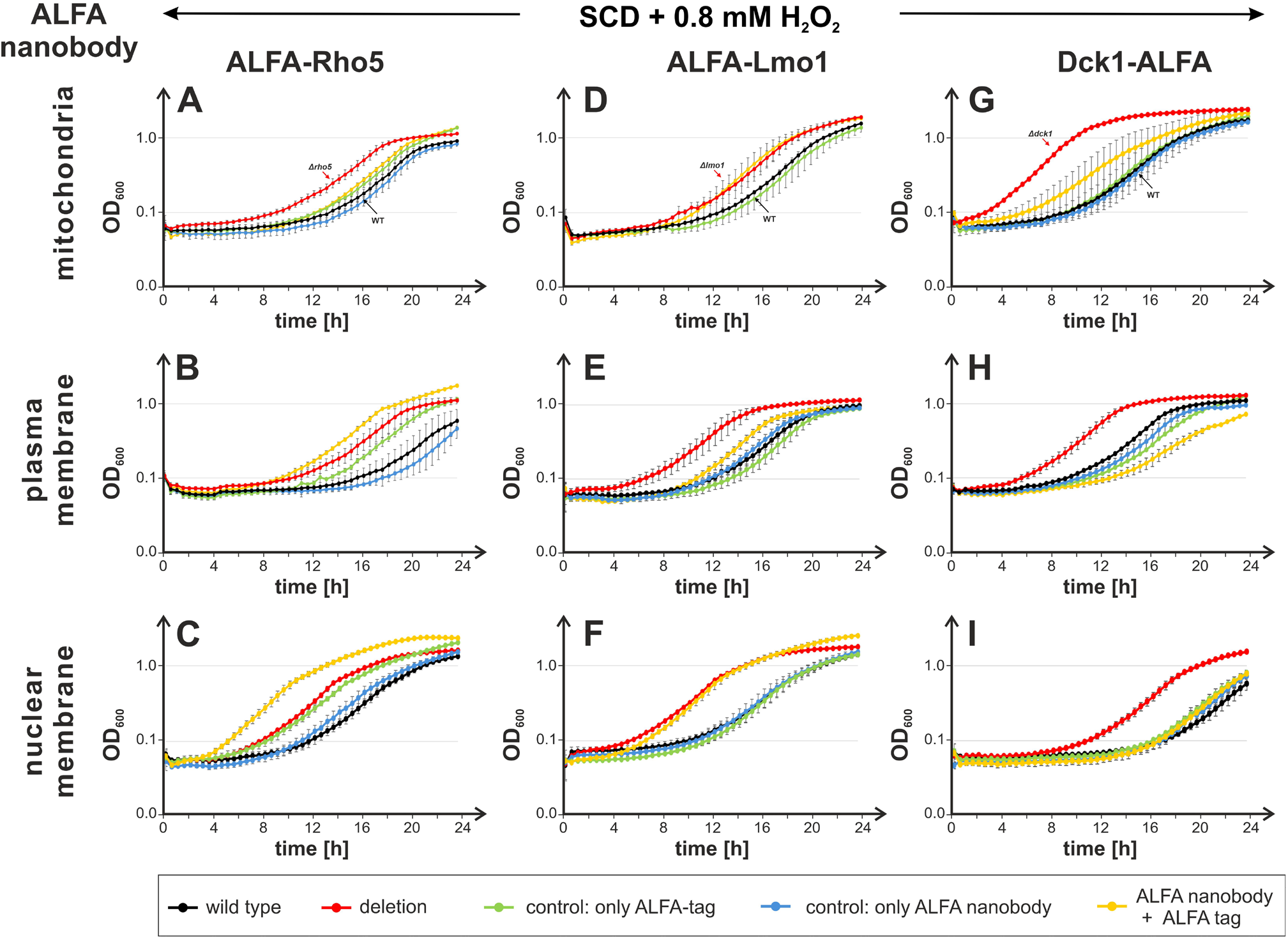
Trapping of Rho5 or its GEF subunits to specific endomembranes impairs growth under oxidative stress. Strains for trapping ALFA-tagged proteins contained nanobody-fusions directed to the outer mitochondrial membrane (ALFAnb-Fis1), the plasma membrane (Mid2-ALFAnb), or the nuclear pores (Nup159-ALFAnb). Growth curves were recorded on synthetic medium with 2% glucose, supplemented with 3 mg/L histidine as required, in the presence of 0.8 mM hydrogen peroxide (controls without hydrogen peroxide showed similar growth for all strains and are shown in supplementary Fig. S1). Error bars give the standard deviation at each time point obtained from at least two biological and two technical replicates from parallel measurements for each row unless indicated otherwise, i.e. two independent isogenic segregants were measured, with two independent inoculates, each. Genotypes of the otherwise isogenic strains are listed in Table 1. The green curves represent the controls for functionality of the three proteins with the ALFA tag; ALFA-Rho5, ALFA-Lmo1 and Dck1-ALFA correspondingly. In blue are the controls for the effect of the ALFA nanobodies; ALFAnb-Fis1, Mid2-ALFAnb and Nup159-ALFAnb respectively. In red are the corresponding deletion strains; *Δrho5, Δlmo1 and Δdck1*. Yellow represents the strains were the corresponding ALFA-tagged Rho5/GEF proteins and ALFA nanobody are present. Strains employed (see Table 1 for genotypes): A) wild type (HOD555-1A and HOD556-2B), *rho5* (HOD555-1D and HOD555-3D), ALFA-Rho5 (HLBO85-7C and HLBO86-4B), ALFAnb-Fis1 (HLBO38-1A, HLBO83-1A and HLBO83-5D), Fis1-Rho5-ALFAtrap (HLBO83-1B, HLBO83-2A and HLBO83-4A). B) wild type (HOD555-1A and HOD556-2B), *rho5* (HOD555-1D and HOD555-3D), ALFA-Rho5 (HLBO83-7D, HLBO85-7C and HLBO86-4B), Mid2-ALFAnb (HLBO85-1D, HLBO85-2C and HLBO85-4C), Mid2-Rho5-ALFAtrap (HLBO85-1A, HLBO85-3B and HLBO85-6C). C) wild type (HD56-5A and HOD556-2B), *rho5* (HOD555-1D and HOD555-3D), ALFA-Rho5 (HLBO83-7D and HLBO85-7C), Nup159-ALFAnb (HD56/AnbH11and HLBO105-1B), Nup159-Rho5-ALFAtrap (HLBO117-1A, HLBO117-1C, HLBO117-6B and HLBO117-6D). D) wild type (HD56-5A and HLBO20-4B), *lmo1* (HOD464-7B and LBO81), ALFA-Lmo1 (LBO309 and LBO310), Fis1-Lmo1-ALFAtrap (HLBO56-2D, HLBO56-5A, HLBO56-11D and HLBO57-5A). E) wild type (HD56-5A and HOD556-2B), *lmo1* (HOD464-7B and LBO81), ALFA-Lmo1 (LBO309 and LBO310), Mid2-ALFAnb (HLBO85-1D and HLBO85-2C), Mid2-Lmo1-ALFAtrap (HLBO84-2B HLBO84-10D). F) wild type (HD56-5A and HOD556-2B), *lmo1* (HOD464-7B **only one**), ALFA-Lmo1 (LBO309 and LBO310), Nup159-ALFAnb (HD56/AnbH11and HLBO105-1B), Nup159-Lmo1-ALFAtrap (HLBO106n-1C and HLBO106n-14C (+Dck1-GFP)). G) wild type (HLBO38-8A **only one**), *dck1* (FSO23-3B and FSO23-6B), Dck1-ALFA (HLBO38-8D and HLBO39-1B), ALFAnb-Fis1 + GFP-Lmo1 (HLBO38-2B and HLBO39-3B), Fis1-Dck1-ALFAtrap (HLBO38-9A and HLBO39-10B). H) wild type (HD56-5A and HOD556-2B), *dck1* (HOD505-1A and HOD505-1C), Dck1-ALFA (HLBO39-1B and HLBO89-2C), Mid2-ALFAnb (HLBO85-1D and HLBO85-2C), Mid2-Dck1-ALFAtrap (HLBO88-10D and HLBO88-12A). I) wild type (HD56-5A and HOD556-2B), *dck1* (HOD505-1A and HOD505-1C), Dck1-ALFA + GFP-Lmo1 (HLBO111-1C **only one**), Nup159-ALFAnb (HD56/AnbH11and HLBO105-1B), Nup159-Dck1-ALFAtrap (HLBO111-8C and HLBO89-1D (with GFP-Lmo1).

Like strains lacking Rho5, lmo1 deletions display a distinct hyper-resistance towards hydrogen peroxide as compared to wild-type strains (Figs. 5D to F). However, trapping this GEF subunit either to the mitochondrial or the nuclear surface fails to restore sensitivity towards this stress, indicating that Lmo1 cannot function when confined to these compartments (Figs. 5D and F). Yet, when trapped at the plasma membrane it appears to be fully functional in this respect (Fig. 5E).

Finally, directing Dck1-ALFA to the different membranes has distinct impacts on the oxidative stress response, as judged from the growth curves recorded in the presence of the stressor (Figs. 5G to I). Thus, the *dck1* deletion shows the expected hyper-resistance towards hydrogen peroxide, whereas the strains recruiting the tagged subunit to the mitochondria (ALFAnb-Fis1^TMD^), are more resistant than wild type, but less than the deletion, indicating a partial functionality Fig. 5G). Trapping to the plasma membrane (Mid2-ALFAnb), or the nuclear membrane (Nup159-ALFAnb) results in growth similar to the wild type. In fact, trapping of Dck1 to the nucleus does not seem to impair function at all (Fig. 5I), while recruitment to the plasma membrane appears to improve GEF function, resembling the hyper-sensitive phenotype of a constitutively active Rho5^G12V^ variant (Fig. 5H; (Sterk et al., 2019)).

## 3. Discussion

The small GTPase Rho5 resides at the plasma membrane under standard growth conditions, but rapidly translocates to the mitochondria upon application of different stresses (Schmitz et al., 2018; Schmitz et al., 2015). Under oxidative stress it has been shown to trigger both mitophagy and apoptosis, as deduced from their strong reduction in *rho5* deletion strains and their hyper-resistance towards agents like hydrogen peroxide (Schmitz et al., 2015). In this work, we addressed the question whether a forced misslocalization of Rho5 or its activating dimeric GEF, composed of Dck1 and Lmo1, mimicked the hyper-resistant phenotype of the deletion mutants, i.e. if the proper translocation to mitochondria under stress is of physiological relevance.

Earlier studies with attempts to direct the three proteins to specific cellular membranes employed either the use of a GFP nanobody fusion to the transmembrane domain of the mitochondrial outer membrane protein Fis1 (Sterk et al., 2019), or a direct fusion of Rho5 to this domain or the cytosolic part of the plasma membrane-borne cell wall integrity sensor Mid2 (Bischof et al., 2022). This approach suffered from the fact that the voluminous GFP-tag impaired the function of the fusion proteins (rendering GFP-Rho5 only partially functional and Lmo1-GFP without function), whereas for some of the direct fusions with Rho5 functionality could not be assessed (Bischof et al., 2022). Therefore, we here decided to employ the ALFA tag, which comprises only 14 amino acid residues, in combination with the respective ALFA nanobody sequence fused to the membrane-targeting proteins. The small tag would be expected to minimize interference with the function of the protein it is attached to, and the nanobody ensures their efficient binding *in vivo* (Gotzke et al., 2019). In general, our findings substantiate these assumptions, as all ALFA-tagged constructs appeared to be functional and were efficiently recruited to the membrane compartments to which the nanobody was attached, as judged from the images of mutual attraction of Lmo1 and Dck1. In addition, working with three candidate proteins, two different tags (ALFA and GFP), and three different membrane compartments, this system allowed a fast and easy combination of the components by genetic crossing and tetrad analysis, as all constructs were obtained in the chromosomes of an isogenic strain series, mostly at their native locus and each with a selectable genetic marker.

Most importantly, we found that the two subunits of the Rho5-GEF, Lmo1 and Dck1, invariably localized together at each membrane that one or the other was trapped at, to the extent that capturing one with the ALFA nanobody the shape of the organelle could be visualized by using a GFP fusion of the other. This strongly suggests that the dimeric GEF spontaneously assembles *in vivo*. We further propose that the rapid translocation of the subunits to mitochondria upon exposure to oxidative stress observed previously in fact occurs in the dimeric form in wild-type cells (Schmitz et al., 2015). Nevertheless, in deletion mutants lacking one of the subunits, the other can still translocate, independently (Bischof et al., 2022).

In contrast to the apparently stable complexes formed by the GEF subunits, Rho5 neither accumulates at the organelles harboring the GEF in the trapping experiments, nor does Rho5 recruit the GEF subunits when it is artificially confined to different membranes, with the possible exception of the slight increase in association with mitochondria without oxidative stress. This may be attributed to the transient nature of their interaction, as the GEF mediates the release of GDP from the GTPase allowing for its substitution by GTP, with its approximately tenfold higher intracellular concentration (Bos et al., 2007). Nucleotide binding is then expected to result in the release of the GTPase from its GEF. Clearly, this would enable Rho5-GTP to traffic within the cell. It is generally assumed that trafficking of Rho-GTPases through the cytosol is mediated in their inactive, GDP-bound conformation by guanine nucleotide dissociation inhibitors (GDI), with Rdi1 being the sole GDI found in yeast (Tiedje et al., 2008). However, the authors did not find any evidence for an interaction of Rdi1 with Rho5, indicating that the latter has other means to traverse the cytosol or remains membrane-bound at all times. As discussed in more detail below, our results favour the diffusion hypothesis rather than the exchange of Rho5 between different membranes through their contact sites. A good candidate for substituting the GDI in masking the hydrophobic lipid anchor may be the long yeast specific extension (LYSE) found in Rho5 between the catalytic core and the C-terminal domains. Deletion of this sequence strongly impairs the translocation of Rho5 to mitochondria under oxidative stress, retaining it at internal structures (Sterk et al., 2019). As the LYSE forms an intrinsically disordered domain, it is tempting to speculate that it could act as kind of an intramolecular GDI (Bischof et al., 2024a).

Besides the interaction studies based on trapping and life-cell fluorescence microscopy, the other important finding regards the physiological role of the distribution of Rho5 and its GEF. In this context, we employed growth under oxidative stress as a functional readout. Previously, *rho5* deletions were shown to be hyper-resistant towards hydrogen peroxide, whereas a constitutively active Rho5^G12V^ variant conferred an increased sensitivity (Schmitz et al., 2015; Singh et al., 2008). Both phenotypes were attributed to the activation of mitophagy and apoptosis by Rho5, triggered by its translocation to the mitochondrial surface (Bischof et al., 2022; Schmitz et al., 2015; Sterk et al., 2019). We reasoned that trapping of the GTPase or its activating GEF at other cellular membranes should interfere with this function and result in a phenotype mimicking that of the respective deletions, i.e. in hyper-resistance towards hydrogen peroxide. Although they reflect a complex relationship, two major conclusions may be drawn from these analyses:

i. For the oxidative stress response, Rho5 appears to be only functional when confined to the mitochondrial surface, but not when trapped at the plasma or nuclear membrane. This implies that it can either be activated by its GEF at this location, or that it can be activated at other sites prior to be trapped by the ALFA nanobody. These trapping data seem somewhat contradictory to previous reports, in which GFP-Rho5 trapped to mitochondria by the respective nanobody appeared to be non-functional, but could be partially recovered if the active GFP-Rho5^G12V^ variant was employed (Sterk et al., 2019). A direct fusion of either Rho5 variant to the transmembrane domain of Fis1 at the mitochondria also lacked the function in oxidative stress response (Bischof et al., 2022). These discrepancies may be explained by the larger tag in the former construct, which rendered it only partially functional, and presumably a lack of function in the direct Fis1 fusion. Thus, we believe our findings herein, with constructs employing the small ALFA tag, to more closely reflect the *in vivo* action of Rho5.
ii. The fact that both Lmo1 and Dck1 appear to be functional when trapped at the plasma membrane, and Lmo1 lacking function at the other two organelles tested, strongly indicates that the GEF can only be activated at this compartment, despite forming a dimer at all intracellular sites, as shown above. In analogy to the human DOCK2/ELMO complex, whose structure has been elucidated with and without the presence of the Rac1 homologue of Rho5, we propose that the GEF first assembles in a closed conformation (Chang et al., 2020; Kukimoto-Niino et al., 2024). Upon interaction with another GTPase, RhoG in the case of the mammalian GEF, it adopts an open conformation, which enables the interaction with nucleotide-bound Rho5. The activatory interaction with Lmo1 may occur exclusively at the plasma membrane with a yet undefined GTPase, explaining the failure of the complex to activate Rho5 when trapped at the mitochondrial or nuclear membrane. Cdc42 may be a good candidate for this interaction, as it acts together with Rho5/Rac1 homologues in other fungi and can partially complement the phenotypes of a *rho5* deletion in the milk yeast *Kluyveromyces lactis* (Hühn et al., 2020; Musielak et al., 2021). This also implies that Rho5-GTP thus produced can translocate to the mitochondrial surface to trigger mitophagy and apoptosis.

The translocation through the cytosol is still a matter of debate, since the sole yeast GDI, Rdi1, apparently does not interact with Rho5 (Tiedje et al., 2008). The extremely fast appearence and even distribution of the GTPase at the mitochondrial surface after application of oxidative stress, i.e. in a matter of seconds (Schmitz et al., 2015), would argue against its diffusion through contact sites between the plasma membrane and mitochondria. This leaves the possibility of another, yet unidentified, yeast protein acting as a chaperone for the hydrophobic lipid anchor, or an intramolecular interaction with the yeast specific extension in Rho5 itself, as suggested above. In the light of the results presented herein, we believe that the latter interaction works with either GDP or GTP bound, thus enabling the translocation of an active Rho5. Clearly, this question merits further investigations.

Our working hypothesis of the GEF being exclusively activated at the plasma membrane is substantiated by the fact that trapping of its two subunits at the mitochondria renders the cells hyper-resistant towards hydrogen peroxide, indicating that Rho5 cannot be suffiently activated. This is also true for misslocalization of Lmo1, but not Dck1, at the nuclear membrane. The observation that a GEF with Dck1 trapped at the nucleus appears to be still functional is difficult to explain. One possibility is that it can acquire Lmo1 that was activated at the plasma membrane at an extent to produce sufficient Rho5-GTP that fulfils its function when reaching the mitochondria. It should also be noted that directing Dck1 to the plasma membrane resulted in an increased sensitivity, reminiscent of the activated Rho5^G12V^ variant (Sterk et al., 2019). Thus, it could reflect a higher amount of activated Rho5 at this site, triggered by the accumulation of the GEF. This is in line with the proposed promotion of an active conformation of the GEF induced by the interaction of Lmo1 with another GTPase at the plasma membrane.

## 4. Material and methods

### 4.1. Yeast strains and media

Isogenic segregants from tetrad analyses of the diploid *S. cerevisiae* strain DHD5 were employed throughout this work (Kirchrath et al., 2000). Names and genotypes of all strains constructed are given in Table 1. For plasmid amplification in *E. coli*, strain DH5α was used (Invitrogen, Karlsruhe, Germany). Standard procedures were followed for genetic manipulations of yeasts and plasmid constructions (Rose et al., 1990). Complete sequences of all plasmids, modified chromosomal loci, and oligonucleotides employed are available upon request. Rich media were based on yeast extract (1% w/v), peptone (2% w/v) and glucose at a concentration of 2% (w/v, YEPD). Synthetic media (SC) contained 0.67% yeast nitrogen base (w/v) supplemented with ammonium sulfate, amino acids and bases as required (Rose et al., 1990), with 2% glucose (w/v, SCD). Histidine concentration was increased from 2 mg/L to 3 mg/L for growth curves, if necessary. *E. coli* cells were grown in LB medium (yeast extract at 0.5% w/v, tryptone at 1% w/v, and sodium chloride at 1% w/v), with the addition of 50 mg/L ampicillin or 25 mg/L kanamycin, as required for plasmid maintenance.

### 4.2. Construction of deletion mutants and strains with tagged genes

Deletion mutants were obtained by one-step gene replacements, using PCR products obtained with primers generating 40–50 bp of homology flanking the genomic target sequences (Rothstein, 1991). Cassettes with selectable genetic markers for deletions were described in (Gueldener et al., 2002), or were modified within the region between the two *loxP* sites to carry the *KlURA3* marker (pJJH1286) or the *KlLEU2* marker (pJJH1287) from *Kluyveromyces lactis*, inserted between the *TEF2* promoter and the *TEF2* terminator from *Ashbya gossypii*. Templates for the mRuby and GFP fluorescence tags were derived from plasmids described in (Kredel et al., 2009; Maeder et al., 2007).

The sequence encoding the ALFA tag (Gotzke et al., 2019) was inserted in the target genes at their native genomic locus in combination with a selective marker to encode either N- or C-terminally tagged proteins, as indicated. Thus, Rho5 and Lmo1 were tagged at the aminoterminus and Dck1 at the carboxyterminus. Expression of the fusion proteins was kept under the control of their native promoters. Sequences of all loci modified can be provided upon request. For the fusion of the ALFA nanobody to Mid2 and Nup159 (PCR-amplified from template plasmids kindly provided by Florian Fröhlich (Osnabrück), the nanobody sequence was amplified by PCR with oligonucleotides generating 40–50 bp of homology flanking the target genomic sequences, producing C-terminal fusion proteins. To direct the ALFA nanobody to the mitochondrial outer membrane, the nanobody sequence was fused to the aminoterminus of the transmembrane domain of Fis1 cloned in an integrative plasmid with *URA3* as a selection marker to yield pJJH3028. The fusion construct was placed under the control of a tailored *PFK2* promoter (Schneider et al., 2018) and thus constitutively expressed after integration into the genomic *ura3-52* allele. For live-cell fluorescence microscopy, functional Dck1– 3myeGFP and GFP-Lmo1 and GFP-Rho5 fusions were encoded at their native genomic loci, using strains described previously (Bischof et al., 2022; Schmitz et al., 2018; Schmitz et al., 2015; Sterk et al., 2019), or otherwise isogenic segregants obtained after crossing and tetrad analysis.

### 4.3. Life-cell fluorescence microscopy

For live-cell fluorescence microscopy cells were grown overnight in synthetic media, diluted to an OD600 of 0.3, and allowed to grow until mid-logarithmic phase with shaking at 30 °C. For fluorescence microscopy a Zeiss Axioplan 2 microscope (Carl Zeiss, Jena, Germany), equipped with a 100x alpha-Plan Fluor objective (NA 1.45) and differential-interference contrast (DIC) was employed. Images were recorded with an ORCA-Flash 4.0 LT Digital CMOS camera C11440 (Hamamatsu Photonics K. K., Japan). Handling of samples and image processing followed previous descriptions (Schmitz et al., 2015). For setup control, the Metamorph v6.2 program (Universal Imaging Corporation,Downingtown, USA) was used. Brightfield images were acquired as single planes using DIC. Fluorescence images were from a single focal point and scaled using Metamorphs scale image command. Images were deconvoluted by either the Huygens Remote Manager v3.7.1 or the Huygens Professional 21.10 program. Processed images were overlaid using Metamorphs overlay images command. Quantification of fluorescence signal at the mitochondria was done manually by counting the cells in blind experiments.

### 4.4. Growth analyses

Growth curves of yeast strains under standard and oxidative stress conditions were recorded with a Varioscan Lux plate reader (ThermoFisher Scientific) as described previously (Sterk et al., 2019). Interval shaking at 30 °C and 1024 rpm was set at cycles of 5 sec on and 7 sec off, using the SkanIt Software 6.0 to control of the plate reader. Path length was equalized to standard photometric measurements by applying a factor of 3.546. Unless specified otherwise, two biological replicates and at least two technical replicates were obtained and averaged for each curve.

## Acknowledgements

We thank Rosaura Rodicio for critical reading of the manuscript and Andrea Murra for technical assistance in the construction of some of the plasmids and fusion alleles.

## Notes

### Competing Interest Statement

The authors have declared no competing interest.

