## Supplementary Figure S1 for "Oxidative stress response mediated by the yeast Rho5 GTPase depends on the proper spatiotemporal distribution of its dimeric GEF"

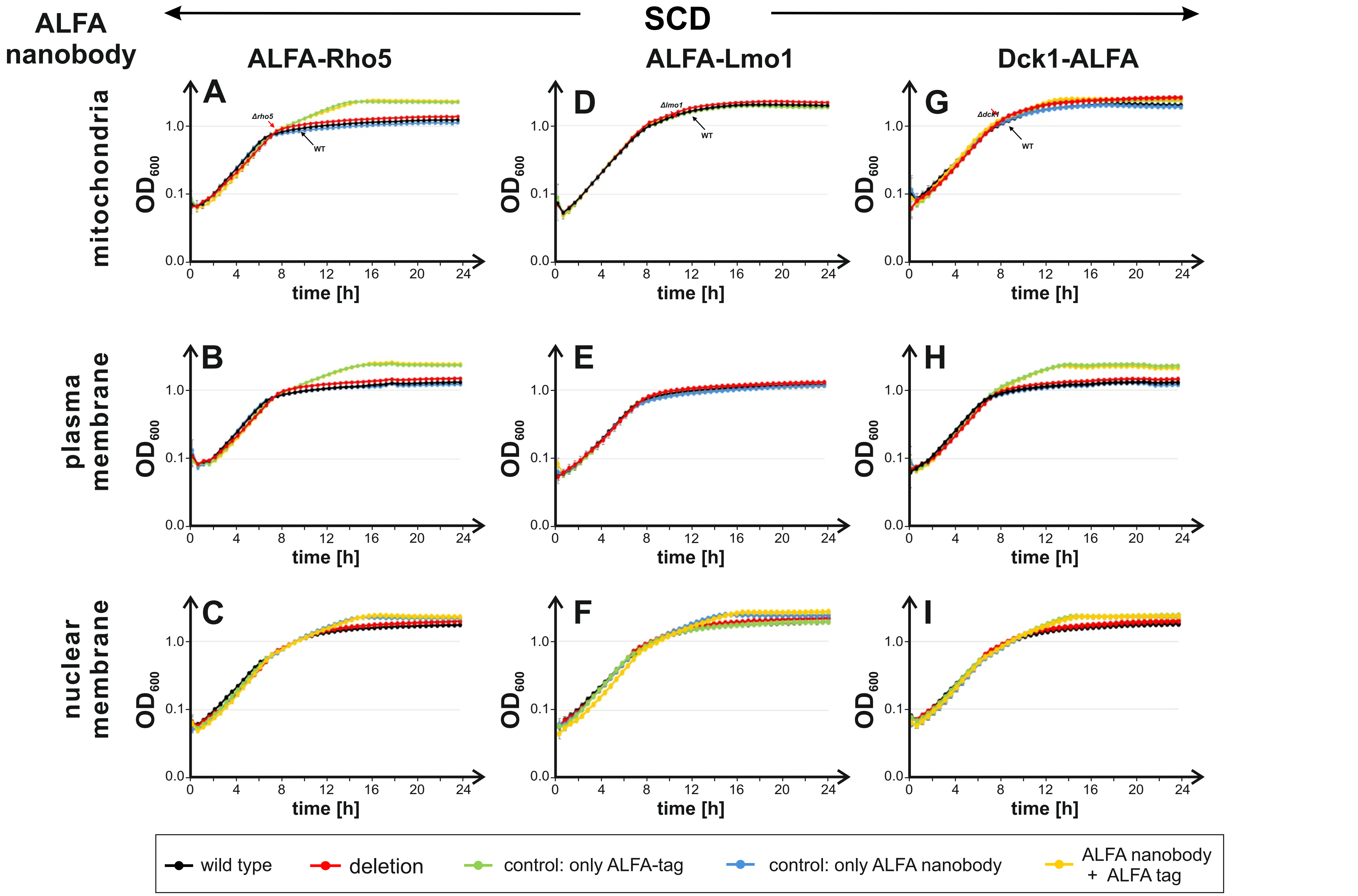


**Figure S1: Trapping of Rho5 or its GEF subunits to specific endomembranes does not impair growth under standard conditions.** Strains for trapping ALFA-tagged proteins contained nanobody-fusions directed to the outer mitochondrial membrane (ALFAnb-Fis1), the plasma membrane (Mid2-ALFAnb), or the nuclear pores (Nup159-ALFAnb). Cells were grown in synthetic complete medium with 2% glucose in the absence of oxidative stress. All other experimental conditions and strains used are the same as described in the legend of Fig. 5 in the main text.
